# Characterization of *P. falciparum* dipeptidyl aminopeptidase 3 specificity reveals structural factors responsible for differences in amino acid preferences between peptide-based substrates and covalent inhibitors

**DOI:** 10.1101/246124

**Authors:** Laura E. de Vries, Mateo I. Sanchez, Katarzyna Groborz, Laurie Kuppens, Marcin Poreba, Christine Lehmann, Fang Yuan, Shirin Arastu-Kapur, Martin Horn, Michael Mares, Matthew Bogyo, Marcin Drag, Edgar Deu

## Abstract

Malarial dipeptidyl aminopeptidases (DPAPs) are cysteine proteases important for parasite development thus making them attractive drug targets. In order to develop inhibitors specific to the parasite enzymes it is necessary to map the determinants of substrate specificity of the parasite enzymes and its mammalian homologue cathepsin C (CatC). Here, we screened peptide-based libraries of substrates and covalent inhibitors to characterize the differences in specificity between parasite DPAPs and CatC, and used this information to develop highly selective DPAP1 and DPAP3 inhibitors. Interestingly, while the primary amino acid specificity of a protease is often used to develop potent inhibitors, we show that equally potent and highly specific inhibitors can be developed based on the sequences of non-optimal peptide substrates. Importantly, analysis of previously published data about the specificity of other proteases also unveiled significant discrepancies in the amino acid preference between substrates and inhibitors. In this article, we also discuss important structural and theoretical reasons that might account for these discrepancies. Overall, this study illustrates that focusing the development of protease inhibitors solely on substrate specificity might overlook important structural features that can be exploited to develop highly potent and selective compounds.

## Introduction

Malaria is a devastating infectious parasitic disease causing nearly half a million deaths every year[1]. Malaria is caused by parasites of the *Plasmodium* genus and is transmitted by *Anopheles* mosquitoes during a blood meal. Within the mosquito midgut, parasites reproduce sexually, multiply, and travel to the salivary glands from where they are transmitted to the human host. Upon infection, the parasites first establish an asymptomatic infection in the liver, followed by an exponential asexual replication in the blood stream, through multiple rounds of red blood cell (RBC) invasion, intracellular replication and egress from infected RBCs, this erythrocytic cycle is responsible for the symptoms and pathology of this disease. Over the last 15 years the world has seen a very significant drop in malaria incidence, mainly due to the global distribution of insecticide-impregnated bed nets and the use of artemisinin-based combination therapies as the standard of care for uncomplicated malaria[2]. However, malaria remains a major global health burden with half of the world population at risk and around 200 million clinical cases per year. Unfortunately, mosquitoes are becoming increasingly resistant to insecticides[3] and artemisinin resistance is on the rise[4], thus making the identification of antimalarial targets and the development of drugs with novel mechanism of action extremely urgent[5].

Dipeptidyl aminopeptidases (DPAPs) are papain-fold cysteine proteases that are expressed at all stages of parasite development[6,7] and might therefore be viable drug targets to treat malaria and prevent its transmission. DPAPs recognize the free N-terminus of protein substrates and cleave N-terminal dipeptides[8,9]. The mammalian homologue cathepsin C (CatC) is the best studied DPAP[10]. In most cells, CatC plays a catabolic lysosomal function. However, in immune cells it is responsible for activating various granule serine proteases involved in the immune response and inflammation such as neutrophil elastase, chymase, granzyme A and B, or cathepsin G[11–14]. Because of its role in activating pro-inflammatory proteases, CatC has been pursued as a potential target for chronic inflammatory diseases[15–17], and phase I clinical trials with CatC inhibitors have been performed by GSK (GSK2793660)[18] and Astrazeneca (AZD7986)[19], thus proving that DPAPs can be targeted with small drug-like molecules.

Three DPAPs are conserved across *Plasmodium* species but very little is known about their molecular functions. In *P. falciparum*, the most virulent *Plasmodium* specie responsible for 90% of malaria mortality, attempts to directly knockout (KO) DPAP1[20] or DPAP3[21] have been unsuccessful, suggesting that they are important for parasite replication. Also, in the *P. berghei* murine model of malaria, KO of DPAP1 or DPAP3 results in a significant decrease in parasite replication[22–24]. DPAP1 localizes mainly in the digestive vacuole[20], an acidic organelle where degradation of haemoglobin takes place. This proteolytic pathway provides a source of amino acids for protein synthesis and liberates space within the RBC for parasites to grow. DPAP1 has been proposed to play an essential role at the bottom of this catabolic pathway[20],[25], however, this function has not yet been confirmed genetically. Previously published inhibition studies suggested that DPAP3 was at the top of the proteolytic cascade that controls parasite egress form iRBCs[26]. However, our recent conditional KO studies have disproven this hypothesis and shown that DPAP3 is instead critical for efficient RBC invasion[21]. Finally, DPAP2 is only expressed in sexual stages and has been shown to be important for gamete egress from iRBCs, thus making it a potential target to block malaria transmission[27],[28]. Overall, a pan-DPAP inhibitor will target the parasite at different stages of development, thus potentially slowing down the emergence of resistance.

A clear understanding of the determinants of substrate specificity of *Plasmodium* DPAPs and CatC will be required in order to develop pan-DPAP inhibitors with minimal off target effects on host CatC, and to design highly specific inhibitors to study the biological function of DPAP1 and DPAP3. A general approach to determine the specificity of proteases upstream of the scissile bond (non-prime pockets) is the use of positional scanning substrate libraries where a fluorophore is conjugated to the C-terminus of a peptide library *via* an amide bond. Proteolytic cleavage of this bond results in a significant increase in fluorescence intensity allowing accurate measurement of substrate turnover. The most common libraries used for this purpose are positional scanning synthetic combinatorial libraries (PS-SCL)[29–31]. PS-SCL are composed of multiple sub-libraries designed to determine the specificity of each non-prime binding pocket in a protease. In each sub-library, the amino acid (AA) at a specific position is varied while a stoichiometric mixture of all natural AAs is used in all other positions. PS-SCL thus provide the substrate specificity at each site in the context of all possible combination of AAs at all other positions. Alternatively, the specificity of a given binding pocket can be determined by varying the identity of the AA at that position while fixing the rest of the peptide to residues known to be recognized by the protease of interest. This approach has been used to fingerprint the specificity of amino exopeptidases such as aminopeptidases[32,33] or DPAPs[34], which only recognize one or two AAs upstream of the scissile bond, respectively. PS-SCL have also been applied to protease inhibitor libraries by replacing the fluorophore with a reversible or irreversible warhead[35]. Optimum substrates and inhibitors are then designed by combining the best residues in each position. Importantly, the recent incorporation of non-natural AAs into these libraries has significantly increased the chemical space that can be explored to characterize the specificity of proteases and has allowed the design of substrates and inhibitors with enhanced selectivity over compounds that contain only natural amino acids[36–38].

Structure-activity relationship (SAR) studies with positional scanning substrate and inhibitor libraries have been performed both on DPAP1 and CatC[25,39,40], but to a much lesser extent on DPAP3[26]. Here, we used libraries of peptide-based substrates and inhibitors to determine the specificity of *P. falciparum* DPAP3 at the P1 and P2 positions. Importantly, the libraries used in this study have been previously screened against DPAP1 and CatC and are therefore ideal to compare the specificities of these three proteases[34]. Our studies show that DPAP3 preferentially cleaves after basic and large aromatic residues (P1 position) and that it prefers substrates having N-terminal aliphatic residues (P2 position). We also identified several non-natural P2 residues that are exclusively recognized by either DPAP1 or DPAP3. By combining the SAR information obtained from these substrate and inhibitor screens we developed specific DPAP1 and DPAP3 inhibitors that remain selective in live parasites. Interestingly, while SAR information obtained from positional scanning substrate libraries is often used to develop potent protease inhibitors[38], we have identified significant differences in specificity between substrates and inhibitors, particularly in the case of DPAP3 and to a lesser extent for CatC and DPAP1. Surprisingly, we also observed significant discrepancies when we compared previously published specificity data about cysteine cathepsins and caspases obtained from PS-SCLs of substrates and inhibitors.

Overall, our study shows that while highly potent inhibitors can be designed based on the sequence of optimal substrates, equally potent and specific inhibitors can be developed using sequences of non-optimal substrates. This work also illustrates how important the choice of warhead is to translate specificity information obtained from positional scanning substrate libraries into potent covalent inhibitors.

## Results

### DPAP3 Substrate Specificity

A positional scanning library of 96 substrates (Fig. 1A), composed of a P1 sub-library of 39 substrates (P2 fixed to Met) and a P2 sub-library of 57 substrates (P1 fixed to homophenylalanine (hPhe)), was screened at 1 μM against recombinant DPAP3 (DPAP3; Fig. 1B-C). The heat map shown in Fig. 1B compares the specificities of DPAP3 with those previously obtained for DPAP1 and CatC[34] at the same substrate concentration. Note that *D*-Phg is the only *D*-AA in P2 that is cleaved by DPAP3, albeit poorly. The remaining 17 substrates containing *D*-AAs in P2 (*D*-hPhe and all natural *D*-AAs except *D*-Cys and *D*-Met) were not cleaved by DPAP3, nor by DPAP1 or CatC. To simplify the Fig., *D*-Phg is the only substrate containing a *D*-AA that is shown in Fig. 1.

**Fig. 1.**
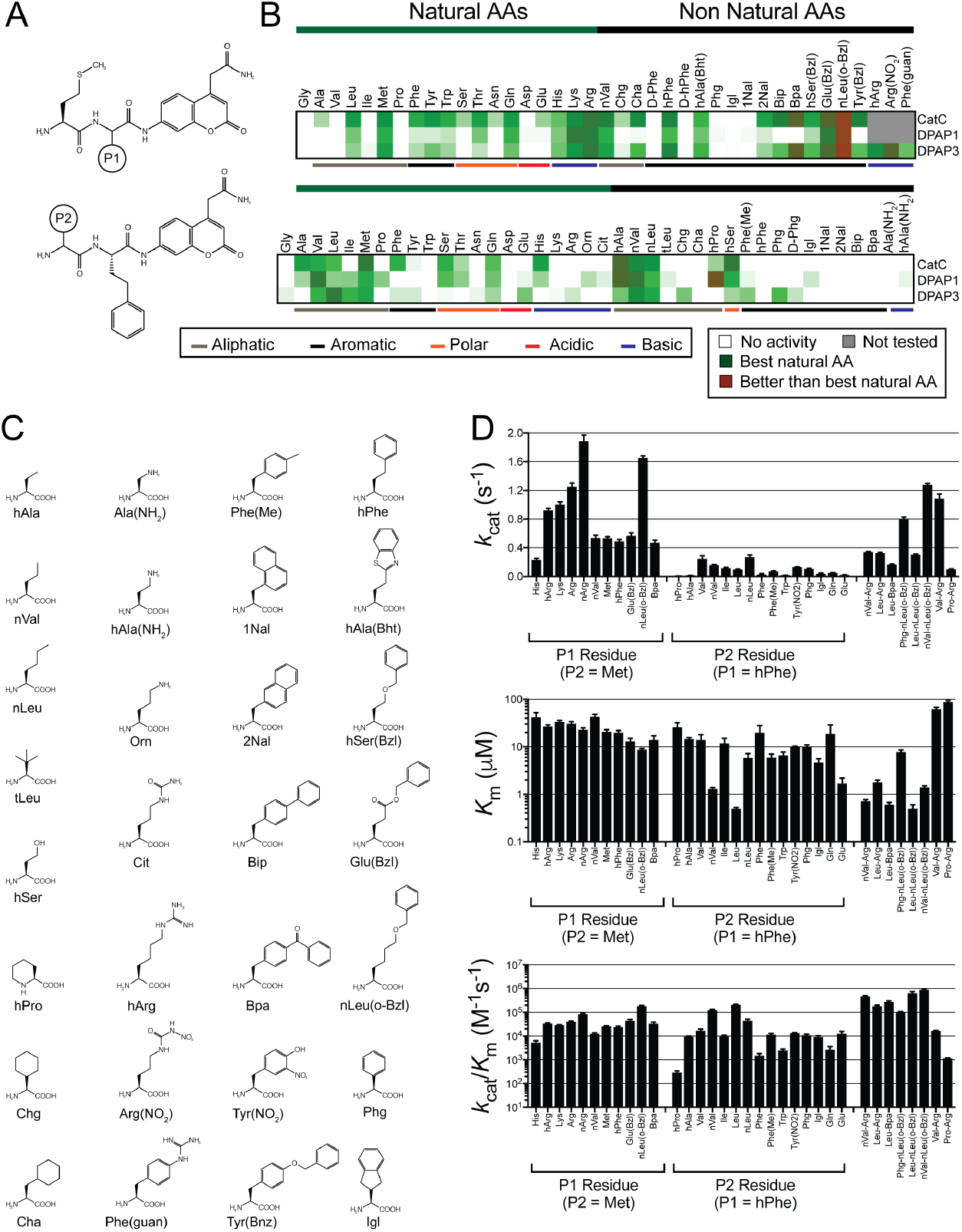
DPAP3 substrate specificity. (**A**) Structure of P1 and P2 substrate libraries. (**B**) Heat map comparing relative turnover rates for the different DPAPs at 1 μM substrate. For each enzyme and substrate turnover rates were normalized relative to the best natural AA (dark green): Arg in P1 for all DPAPs; Met, Val, and Leu in P2 for CatC, DPAP1, and DPAP3, respectively. Red indicates substrates that are turned over better than the best natural AA. White represent no activity, and grey substrates that were only tested on DPAP3. (**C**) Structure of non-natural AAs used in the substrate library. (**D**) Steady-state Michaelis Menten parameters for DPAP3 determined for selected substrates.

All three DPAPs show broad and similar P1 specificity, which is not surprising because in clan CA proteases the P1 residue side chain is solvent exposed. For all DPAPs, a general preference for long basic, aliphatic, and aromatic P1 residues was observed—basic: Lys, Arg, homoarginine (hArg), and nitroarginine (Arg(NO_2_)); aliphatic: Met, norvaline (nVal) and Leu; aromatic: hPhe, (4-benzothiazol-2-yl)homoalanine (hAla (Bht)), 6-benzyloxynorleucine (nLeu(o-Bzl)), glutamic acid benzyl ester (Glu(Bzl)), and homoserine-*O*-benzyl (hSer(Bzl)) —. Interestingly, DPAP1 differs from CatC and DPAP3 in that it does not accept large hydrophobic groups such as cyclohexylalanine (Cha), 2-naphthalene (2Nal), biphenylalanine (Bip), 4-benzoyl-phenylalanine (Bpa), tyrosine-*O*-benzoyl (Tyr(Bzl)), or 4-guanidino-phenylalanine (Phe(guan)) in P1. Unfortunately, we could not identify any P1 residue that was recognized by DPAP1 and/or DPAP3 but not by CatC.

Clear differences in specificity were observed between the three DPAPs at the P2 position (Fig. 1B). DPAP3 seems to have a narrower P2 specificity for natural AAs than DPAP1 or CatC, probably reflecting its more specific biological function in RBC invasion compared to the catabolic functions of DPAP1 and CatC. Both DPAP1 and DPAP3 have a strong preference for long aliphatic residues such as Leu, Ile, norleucine (nLeu), Met, or norvaline (nVal). Some non-natural AAs seem to be exclusively cleaved by DPAP3 such as cyclohexylglycine (Chg), Phg, or 4-methyl-phenylalanine (Phe(Me)). Surprisingly, Phe(Me) is the only substrate in the library with an aromatic P2 residue that is accepted by DPAP3 even though vinyl sulfone (VS) inhibitors with P2 aromatic residues such as Tyr, Trp, or nitrotyrosine (Tyr(NO_2_)) have been shown to be potent DPAP3 inhibitors[21,26]. Interestingly, an Ile in P2 is efficiently cleaved by both *Plasmodium* DPAPs but not by CatC.

### Development of optimum DPAP3 substrates

To determine how P1 and P2 side chains influence *k*_cat_ and *K*_m_ values for DPAP3, we performed Michaelis Menten studies on selected substrates from the P1 and P2 libraries. In addition, we synthesized a series of substrates that combine optimal natural and non-natural AAs for DPAP3: Arg, hPhe, nLeu(o-Bzl), and Bpa in P1, and Leu, Val, nVal, and Phg in P2. We also tested substrates predicted to be DPAP1-selective (Pro-Arg-ACC and hPro-hPhe-ACC) against DPAP3. Finally, because we were surprised by the lack of activity observed for substrates with aromatic P2 residues, we measured Michaelis Menten parameters for Phe-Arg-ACC, Trp-hPhe-ACC, and Tyr(NO_2_)-hPhe-ACC. The sequence of the last substrate is based on the structure of SAK1 (Tyr(NO_2_)-hPhe-VS), which is the most potent DPAP3 inhibitor identified so far (see below). Table 1 and Fig. 1D report the Michaelis Menten parameters determined for all these substrates (Michaelis Menten curves are shown in Fig. S1).

**Table 1.** Steady-state Michaelis Menten parameters for DPAP3.

| **Table1.** Steady-state Michaelis Menten parameters for DPAP3. |  |  |  |  |
| --- | --- | --- | --- | --- |
| Substrate | N | $k_{cat}$<br>(s<sup>-1</sup>) | $K_m$<br>(μM) | $k_{cat}/K_m \cdot 10^{-3}$ (M<sup>-1</sup>s<sup>-1</sup>) |
| Met-His-ACC | 4 | 0.23 ± 0.02 | 42 ± 10 | 5.4 ± 1 |
| Met-nVal-ACC | 4 | 0.54 ± 0.04 | 43 ± 5 | 13 ± 1 |
| Met-hPhe-ACC | 4 | 0.5 ± 0.02 | 20 ± 2 | 24 ± 2 |
| Met-Met-ACC | 4 | 0.54 ± 0.02 | 21 ± 2 | 26 ± 1 |
| Met-Lys-ACC | 4 | 1.00 ± 0.04 | 34 ± 2 | 29 ± 1 |
| Met-Bpa-ACC | 3 | 0.48 ± 0.04 | 14 ± 3 | 34 ± 4 |
| Met-hArg-ACC | 4 | 0.92 ± 0.02 | 27 ± 2 | 35 ± 1 |
| Met-Arg-ACC | 4 | 1.26 ± 0.06 | 31 ± 3 | 41 ± 2 |
| Met-Glu(Bzl)-ACC | 4 | 0.56 ± 0.04 | 13 ± 2 | 46 ± 6 |
| Met-Arg(NO<sub>2</sub>)-ACC | 4 | 1.90 ± 0.08 | 23 ± 2 | 84 ± 4 |
| Met-nLeu(o-Bzl)-ACC | 10 | 1.66 ± 0.02 | 8.8 ± 0.4 | 186 ± 6 |
| hPro-hPhe-ACC<sup>1</sup> | 3 | 0.007 ± 0.001 | 26 ± 6 | 0.30 ± 0.04 |
| Trp-hPhe-ACC | 3 | 0.017 ± 0.002 | 7 ± 1 | 2.5 ± 0.3 |
| Phe-hPhe-ACC | 3 | 0.032 ± 0.008 | 20 ± 8 | 1.6 ± 0.2 |
| Igl-hPhe-ACC | 3 | 0.044 ± 0.004 | 4.7 ± 0.9 | 9.4 ± 1 |
| hAla-hPhe-ACC | 3 | 0.0014 ± 0.002 | 14.7 ± 0.8 | 9.8 ± 1 |
| Ile-hPhe-ACC | 4 | 0.12 ± 0.01 | 12 ± 3 | 10 ± 1 |
| Phg-hPhe-ACC | 4 | 0.110 ± 0.004 | 10 ± 1 | 11 ± 1 |
| Phe(Me)-hPhe-ACC | 3 | 0.072 ± 0.004 | 6 ± 1 | 12 ± 1 |
| Tyr(NO<sub>2</sub>)-hPhe-ACC | 3 | 0.134 ± 0.002 | 9.9 ± 0.4 | 13.6 ± 0.4 |
| Gln-hPhe-ACC | 3 | 0.052 ± 0.001 | 19 ± 10 | 2.7 ± 0.9 |
| Glu-hPhe-ACC | 3 | 0.024 ± 0.002 | 1.7 ± 0.5 | 13 ± 3 |
| Val-hPhe-ACC | 3 | 0.24 ± 0.04 | 14 ± 4 | 18 ± 2 |
| nLeu-hPhe-ACC | 3 | 0.28 ± 0.02 | 5.9 ± 1.3 | 46 ± 6 |
| nVal-hPhe-ACC | 4 | 0.164 ± 0.005 | 1.30 ± 0.08 | 128 ± 6 |
| Leu-hPhe-ACC | 3 | 0.100 ± 0.002 | 0.50 ± 0.03 | 210 ± 10 |
| Phg-nLeu(o-Bzl)-ACC | 8 | 0.80 ± 0.02 | 7.8 ± 0.8 | 102 ± 6 |
| Leu-nLeu(o-Bzl)-ACC | 5 | 0.308 ± 0.006 | 0.5 ± 0.1 | 640 ± 100 |
| nVal-nLeu(o-Bzl)-ACC | 8 | 1.28 ± 0.02 | 1.4 ± 0.1 | 920 ± 60 |
| Leu-Bpa-ACC | 4 | 0.170 ± 0.004 | 0.61 ± 0.06 | 280 ± 20 |
| Phe-Arg-ACC | 4 | 0.018 ± 0.006 | 20 ± 10 | 0.9 ± 0.2 |
| Pro-Arg-AMC<sup>2</sup> | 3 | 0.102 ± 0.004 | 88 ± 5 | 1.16 ± 0.04 |
| Val-Arg-ACC<sup>3</sup> | 3 | 1.08 ± 0.06 | 63 ± 5 | 17.0 ± 0.4 |
| Leu-Arg-ACC | 8 | 0.330 ± 0.006 | 1.8 ± 0.2 | 185 ± 15 |
| nVal-Arg-ACC | 8 | 0.346 ± 0.004 | 0.72 ± 0.05 | 480 ± 30 |
| *Phe-Arg-βNA* | 4 | 0.96 ± 0.04 | 51 ± 5 | 19 ± 1 |

N is the number of replicates.
<sup>1</sup>For DPAP1, $k_{cat} = 0.22 \pm 0.004$ s<sup>-1</sup>, $K_m = 0.67 \pm 0.14$ μM and $k_{cat}/K_m = 320,000 \pm 50,000$ M<sup>-1</sup>s<sup>-1</sup>; for CatC, $k_{cat} = 10.8 \pm 0.2$ s<sup>-1</sup>, $K_m = 91 \pm 5$ μM and $k_{cat}/K_m = 116,000 \pm 11,000$ M<sup>-1</sup>s<sup>-1</sup>. [34]
<sup>2</sup>For DPAP1, $k_{cat} = 6.2 \pm 0.4$ s<sup>-1</sup>, $K_m = 84 \pm 9$ μM and $k_{cat}/K_m = 74,000 \pm 2000$ M<sup>-1</sup>s<sup>-1</sup>; for CatC, $k_{cat} = 490 \pm 10$ s<sup>-1</sup>, $K_m = 130 \pm 10$ μM and $k_{cat}/K_m = 3,600,000 \pm 300,000$ M<sup>-1</sup>s<sup>-1</sup>. [40]
<sup>3</sup>For DPAP1, $k_{cat} = 3.5 \pm 0.1$ s<sup>-1</sup>, $K_m = 21 \pm 2$ μM and $k_{cat}/K_m = 170,000 \pm 10,000$ M<sup>-1</sup>s<sup>-1</sup>; for CatC, $k_{cat} = 180 \pm 10$ s<sup>-1</sup>, $K_m = 51 \pm 8$ μM and $k_{cat}/K_m = 3,600,000 \pm 300,000$ M<sup>-1</sup>s<sup>-1</sup>. [40]

P1 residues have a significant influence in *k*_cat_, with nLeu(o-Bzl) and positively charged residues having the highest values. A positive charge on the δ position (Arg(NO_2_) and Arg) is favoured over the e position (Lys and hArg). Elongated aliphatic and hydrophobic residues in P1 decrease *K*_m_, especially when aromatic groups are distant from the peptide backbone. This is evident by the decreasing *K*_m_ values between nVal, Met, hPhe, Bpa, Glu(Bzl), and nLeu(o-Bzl). This tendency was also observed for CatC and DPAP1, and might suggest the presence of a distal binding pocket (Fig. 1B), potentially an exosite, since P1 residues are usually solvent exposed in clan CA proteases.

P2 residues have a bigger influence on *K*_m_ than P1, with Leu and nVal being optimal P2 residues for DPAP3. Beta branched residues are not optimal for DPAP3 as can be observed by an increase in *K*_m_ between nVal and Val or nLeu and Ile. However, the g-branched AA Leu has the lowest *K*_m_. Substrates with aliphatic P2 side chains that extend to the δ position (Met and nLeu) result in higher *K*_m_ values than slightly shorter ones (Leu and nVal) but also higher *k*_cat_ values. Overall, combining optimal P1 (nLeu(o-Bzl) and Arg) and P2 (nVal and Leu) residues results in improved *k*_cat_/*K*_m_ values (Table 1).

Interestingly, although substrates with Phg and indanyl-glycine (Igl) in P2, or Bpa in P1, are not the preferred AAs at these positions, these non-natural residues are structurally very different from natural AAs and are turned over quite efficiently by DPAP3 when combined with optimal P1 or P2 residues, respectively, i.e. Leu-Bpa-ACC or Phg-nLeu(o-Bzl)-ACC. Finally, the optimal substrate for DPAP1, hPro-hPhe-ACC[34], is very poorly turned over by DPAP3 (> 200-fold difference in *k*_cat_/*K*_m_). We think that substrates containing these non-natural AAs could be used as specific tools to measure DPAP1 or DPAP3 activity in biological samples, i.e. parasite lysates or live parasites, an application we are currently investigating.

Finally, our studies show that substrates with aromatic P2 residues are poorly cleaved by DPAP3 compared to optimal substrates, i.e. 100 to 1000-fold lower *k*_cat_/*K*_m_. This is surprising since vinyl sulfone inhibitors containing aromatic P2 residues such as Tyr(NO_2_) or Trp, are potent and selective DPAP3 inhibitors[21,26]. Because these two AA side chains have fluorogenic properties, we investigated whether the low turnover rate measured for Tyr(NO_2_)-hPhe-ACC and Trp-hPhe-ACC might be due to quenching effects. The emission of free ACC (0, 1, or 5 μM) in assay buffer was measured in the presence of 0-100 μM of these substrates (Fig. 2A-B). No significant decrease the ACC emission signal was observed even when substrates were present in 100-fold excess, thus indicating that the low turnover rates measured for these substrates are not due to quenching effects.

**Fig. 2.**
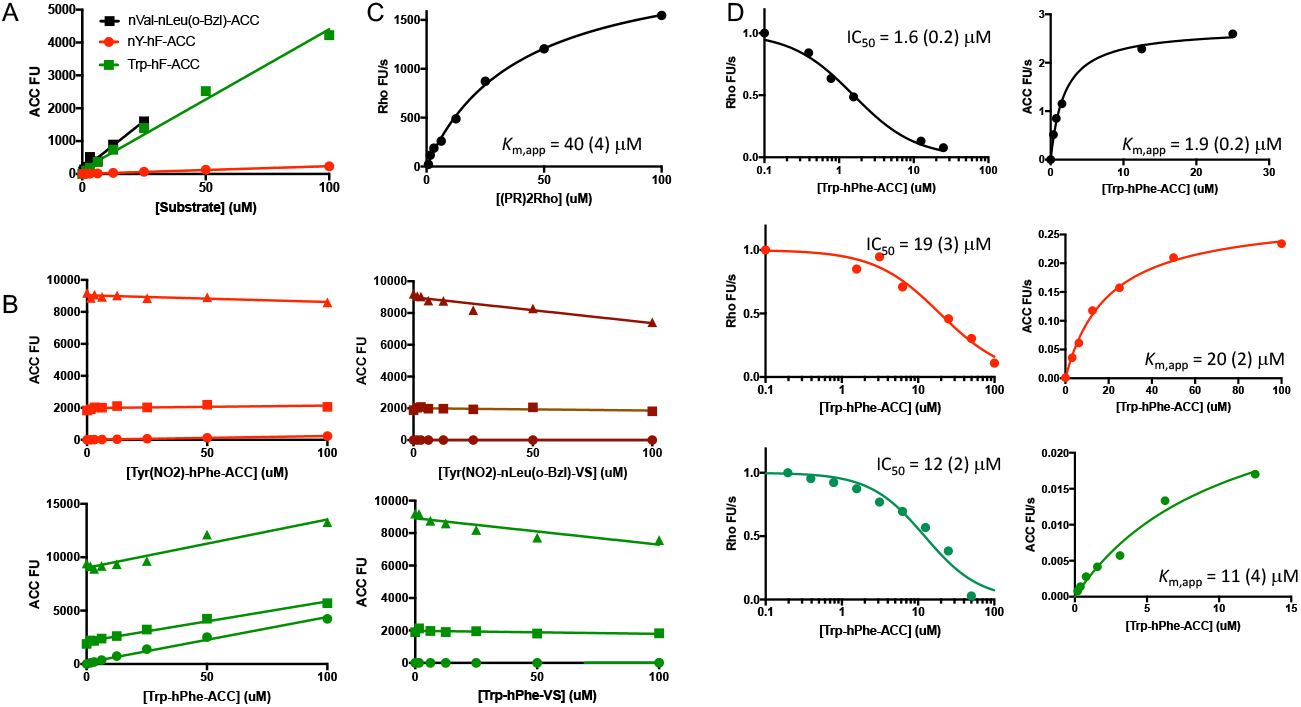
Substrate quenching and substrate competition studies. (**A**) Background fluorescence signal of selected substrates measured in assay buffer. Note that Tyr(NO_2_)-hPhe-ACC shows a very low level of background fluorescence compared to Trp-hPhe-ACC or nVal-nLeu(o-Bzl)-ACC (Fig. 3A) likely indicating intramolecular quenching between ACC and Tyr(NO_2_). (**B**) ACC fluorescence signal measured at increasing concentrations of the indicated substrates and inhibitors in the presence at 0 (circles), 1 (squares) or 5 (triangles) μM of free ACC. (**C**) (PR)_2_Rho turnover by rDPAP3. (**D**) Turnover rates of (PR)_2_Rho (left graphs) and ACC substrates (right graphs) measured at 40 μM of (PR)_2_Rho and increasing concentrations of ACC substrates. *K*_m,app_ and IC_50_ values are indicated in each graph.

As an alternative method to confirm that Tyr(NO_2_)-hPhe-ACC and Trp-hPhe-ACC bind relatively poorly to DPAP3, we performed substrate competition assays using the (PR)_2_Rho substrate (*λ*_ex_ = 492 nm, *λ*_em_ = 523 nm), which emits at much higher wavelengths than ACC (*λ*_ex_ = 355 nm, *λ*_ex_ = 460 nm), and thus allowing us to simultaneously measure the turnover of (PR)_2_Rho and ACC substrates without quenching interference. (PR)_2_Rho was initially designed as a DPAP1 specific substrate to directly measure the activity of this protease in crude parasite extracts[41], but it is also cleaved by DPAP3 with a *K*_m,app_ of 40 μM (Fig. 3C). This substrate is cleaved twice by DPAPs, releasing two Pro-Arg dipeptides and the rhodamine 110 fluorophore. In this assay, we simultaneously measured inhibition of (PR)_2_Rho turnover by ACC substrates (IC_50_ values) as well as the apparent *K*_m_ values of these ACC substrates in the presence of 40 μM (PR)_2_Rho (*K*_m,app_ values). As shown in Fig. 2D, the IC_50_ and *K*_m,app_ values obtained are within experimental error and, as expected, slightly higher than the *K*_m_ values reported in Table 1 due to the substrate competition effect.

**Fig. 3.**
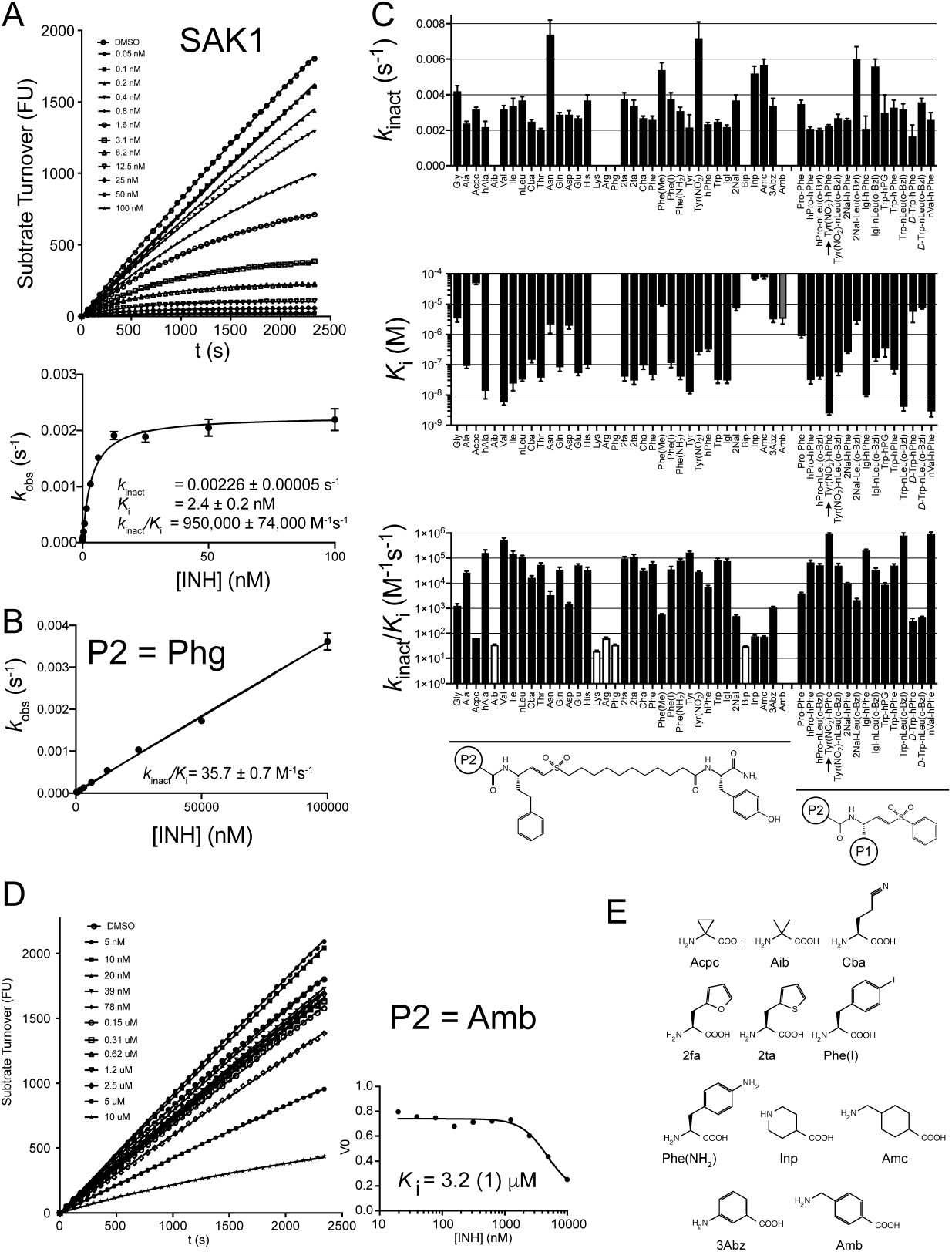
DPAP3 inhibitor specificity. (**A**) Representative data showing time-dependent inhibition of DPAP3 by SAK1 (Tyr(NO_2_)-hPhe-VS). Each progress curve (FU vs. time) was fitted to Eq. 5 to obtain *k*_obs_ values (top graph). These were then fitted to Eqs. 6 and 7 to obtain *k*_inact_, *K*_i_ and *k*_inact_/*K*_i_. (**B**) Example of an inhibitor where no inhibitor saturation was observed (white bars in C). In this case, *k*obs values were fitted to a linear model to obtain *k*_inact_/*K*_i_. (**C**) Inhibition constants determined for DPAP3. The general structure of the inhibitors is shown below the graphs. White bars correspond to inhibitors for which only *k*_inact_/*K*_i_ values could be determined. The grey bar corresponds to the only inhibitor that showed a reversible mechanism of inhibition, i.e. only a *K*_i_ value could be determined. The inhibitor corresponding to SAK1 is indicated with an arrow. (**D**) No time-dependence inhibition of DPAP3 was observed with an inhibitor with a P2 Amb. Initial turnover rates (*V*_0_) were fitted to a reversible binding model to obtain *K*_i_. (**E**) Structure of non-natural AAs present in the inhibitor library but not in the substrate library.

Overall, the lack of quenching effect between Trp or Tyr(NO_2_) and ACC, and the good agreement between IC_50_ and *K*_m,app_ measured under substrate competition conditions indicate that the Michaelis Menten parameters reported in Table 1 are accurate and that substrates containing aromatic P2 AAs are indeed relatively poor DPAP3 substrates compared to those containing optimal aliphatic P2 residues. These results raise the question of why vinyl sulfone inhibitors with aromatic P2 residues are among the most potent DPAP3 inhibitors identified so far. To better understand this discrepancy, we measured the kinetics of inactivation of DPAP3 by the previously published vinyl sulfone inhibitor library[25,26].

### DPAP3 Inhibitor Specificity

Time- and concentration-dependent inactivation of DPAP3 by the P2 library of vinyl sulfone inhibitors (P1 fixed to hPhe) was measured using a continuous assay at 2.2 μM of Met-nLeu(o-Bzl)-ACC (0.25 x *K*_m_). For most compounds, the mechanism of inhibition was consistent with a two-step irreversible inhibition model (Eq. 1, Fig. 3A).

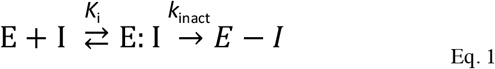

*K*_i_ is the inhibition equilibrium constant, and *k*_inact_ the rate of covalent modification of the catalytic Cys.

For a few inhibitors only *k*_inact_/*K*_i_ values could be obtained, i.e. no saturation was achieved in the *k*_obs_ *vs*. [I] graph (Fig. 3B). The inhibition constants are reported in Fig. 3C and Table 2, and the curve fits shown in Fig. S2. Only one inhibitor, the one with an amino-methyl-benzyl (Amb) group in P2, was not able to inhibit DPAP3 in a time-dependent manner under our assay conditions (Fig. 3D). This is probably due to the fact that this extended and rigid P2 AA (Fig. 3E) might prevent proper positioning of the vinyl sulfone group into the active site of DPAP3 to allow covalent modification of the catalytic Cys. For this compound, a *K*_i_ value for reversible inhibition was measured.

**Table 2.** DPAP3, DPAP1 and CatC inhibition constants for the vinyl sulfone library.

| P2 | DPAP3 |  |  | DPAP1 |  |  | CatC |  |  |
| --- | --- | --- | --- | --- | --- | --- | --- | --- | --- |
| | $k_{\text{inact}} \cdot 10^{-3}$<br>(s<sup>-1</sup>) | $K_i$<br>(nM) | $k_{\text{inact}}/K_i \cdot 10^{-3}$<br>(M<sup>-1</sup>s<sup>-1</sup>) | $k_{\text{inact}} \cdot 10^{-3}$<br>(s<sup>-1</sup>) | $K_i$<br>(nM) | $k_{\text{inact}}/K_i \cdot 10^{-3}$<br>(M<sup>-1</sup>s<sup>-1</sup>) | $k_{\text{inact}} \cdot 10^{-3}$<br>(s<sup>-1</sup>) | $K_i$<br>(nM) | $k_{\text{inact}}/K_i \cdot 10^{-3}$<br>(M<sup>-1</sup>s<sup>-1</sup>) |
| Gly | 4.2 ± 0.3 | 3,300 ± 750 | 1.30 ± 0.25 | 3.0 ± 0.2 | 3,300 ± 750 | 0.9 ± 0.1 | 3.0 ± 0.2 | 70 ± 9 | 43 ± 4 |
| Ala | 2.4 ± 0.1 | 88 ± 13 | 27 ± 3 | 3.03 ± 0.09 | 90 ± 7 | 34 ± 2 | 3.00 ± 0.06 | 9.7 ± 0.4 | 310 ± 8 |
| Acpc | 3.2 ± 0.1 | 47,000 ± 3,000 | 0.0677 ± 0.0001 | 6.2 ± 0.3 | 49,000 ± 4,000 | 0.127 ± 0.006 | 1.03 ± 0.06 | 3,200 ± 700 | 0.33 ± 0.06 |
| hAla | 2.2 ± 0.3 | 13.5 ± 6 | 166 ± 50 | 5.2 ± 0.7 | 19 ± 4 | 267 ± 26 | 2.9 ± 0.2 | 5.9 ± 0.9 | 500 ± 50 |
| Aib | N.S. | N.S. | 0.0363 ± 0.0006 | 8.9 ± 0.5 | 18,000 ± 2,000 | 0.49 ± 0.03 | 1.20 ± 0.07 | 2,100 ± 300 | 0.57 ± 0.06 |
| Val | 3.2 ± 0.2 | 5.7 ± 1.1 | 554 ± 84 | 3.6 ± 0.2 | 2.6 ± 0.4 | 1,380 ± 140 | 2.9 ± 0.1 | 14 ± 2 | 200 ± 15 |
| Ile | 3.4 ± 0.4 | 23 ± 9 | 148 ± 45 | 3.2 ± 0.1 | 92 ± 8 | 34 ± 3 | 3.7 ± 0.8 | 1,300 ± 400 | 2.9 ± 0.3 |
| nLeu | 3.7 ± 0.2 | 31 ± 3 | 120 ± 7.5 | 4.4 ± 0.3 | 14 ± 2 | 305 ± 24 | 2.40 ± 0.05 | 7.0 ± 0.5 | 340 ± 20 |
| Cba | 2.5 ± 0.1 | 140 ± 25 | 17 ± 2.5 | 5.4 ± 0.4 | 13,000 ± 2,000 | 0.41 ± 0.03 | 1.29 ± 0.03 | 2,400 ± 140 | 0.53 ± 0.02 |
| Thr | 2.0 ± 0.1 | 36 ± 8 | 55 ± 10 | 4.4 ± 0.2 | 112 ± 10 | 39 ± 2 | 2.38 ± 0.03 | 65 ± 3 | 36 ± 1 |
| Asn | 7.4 ± 0.8 | 2,100 ± 1,000 | 3.5 ± 1.4 | 9.9 ± 0.5 | 10,500 ± 750 | 0.95 ± 0.02 | 2.40 ± 0.07 | 260 ± 30 | 9.2 ± 0.7 |
| Gln | 2.9 ± 0.2 | 80 ± 20 | 35 ± 8 | 4.7 ± 0.2 | 50 ± 3 | 93 ± 3 | 2.3 ± 0.1 | 15 ± 3 | 160 ± 25 |
| Asp | 2.9 ± 0.2 | 1,900 ± 400 | 1.5 ± 0.2 | 5.4 ± 0.2 | 2,100 ± 200 | 2.6 ± 0.10 | 3.0 ± 0.1 | 2,200 ± 200 | 1.4 ± 0.2 |
| Glu | 2.7 ± 0.1 | 51 ± 8 | 53 ± 6 | 5.0 ± 0.4 | 650 ± 100 | 7.6 ± 0.75 | 2.6 ± 0.1 | 140 ± 20 | 18 ± 2 |
| His | 3.7 ± 0.3 | 100 ± 25 | 35.5 ± 7.5 | 4.3 ± 0.3 | 150 ± 20 | 29 ± 3 | 2.1 ± 0.1 | 0.6 ± 0.1 | 3,350 ± 500 |
| Lys | N.S. | N.S. | 0.020 ± 0.001 | N.S. | N.S. | 0.081 ± 0.004 | 1.49 ± 0.05 | 3,600 ± 200 | 0.42 ± 0.02 |
| Arg | N.S. | N.S. | 0.065 ± 0.006 | N.S. | N.S. | 0.084 ± 0.002 | 4.0 ± 0.5 | 11,000 ± 2,000 | 0.38 ± 0.02 |
| Phg | N.S. | N.S. | 0.0357 ± 0.0007 | 8.3 ± 0.5 | 7,000 ± 650 | 1.19 ± 0.05 | 2.5 ± 0.2 | 86 ± 20 | 29 ± 5 |
| 2fa | 3.8 ± 0.3 | 39 ± 10 | 97 ± 18 | 5.2 ± 0.4 | 11 ± 3 | 470 ± 90 | 3.7 ± 0.2 | 2.1 ± 0.2 | 1,720 ± 90 |
| 2ta | 3.4 ± 0.3 | 29 ± 7 | 118 ± 23 | 3.8 ± 0.3 | 82 ± 20 | 47 ± 7 | 3.3 ± 0.3 | 6.5 ± 1 | 510 ± 55 |
| Cha | 2.7 ± 0.1 | 90 ± 20 | 31.6 ± 5.5 | N.S. | N.S. | 0.020 ± 0.001 | 2.3 ± 0.1 | 300 ± 40 | 7.7 ± 0.8 |
| Phe | 2.6 ± 0.2 | 45 ± 13 | 60 ± 15 | N.S. | N.S. | 0.40 ± 0.01 | 2.18 ± 0.08 | 1.4 ± 0.2 | 1,600 ± 200 |
| Phe(Me) | 5.4 ± 0.4 | 9,200 ± 300 | 0.580 ± 0.025 | N.S. | N.S. | 0.314 ± 0.005 | 2.8 ± 0.2 | 1,300 ± 200 | 2.2 ± 0.3 |
| Phe(I) | 3.8 ± 0.3 | 110 ± 30 | 36 ± 10 | 7.0 ± 0.3 | 47,000 ± 3,500 | 0.151 ± 0.006 | 1.02 ± 0.06 | 185 ± 40 | 5.5 ± 0.9 |
| Phe(NH2) | 3.1 ± 0.2 | 39 ± 7 | 80 ± 12 | 5.0 ± 0.4 | 22,000 ± 2500 | 0.23 ± 0.01 | 1.01 ± 0.02 | 5.3 ± 0.3 | 191 ± 9 |
| Tyr | 2.18 ± 0.07 | 12.7 ± 1.5 | 172 ± 16 | N.S. | N.S. | 0.310 ± 0.005 | 2.10 ± 0.06 | 11.5 ± 0.9 | 182 ± 10 |
| Tyr(NO<sub>2</sub>) | 7.2 ± 0.9 | 250 ± 40 | 28.8 ± 0.8 | N.S. | N.S. | 0.023 ± 0.002 | N.S. | N.S. | 0.147 ± 0.008 |
| hPhe | 2.36 ± 0.08 | 310 ± 30 | 7.6 ± 0.7 | N.S. | N.S. | 0.029 ± 0.002 | N.S. | N.S. | 0.103 ± 0.005 |
| Trp | 2.5 ± 0.1 | 30 ± 6 | 84 ± 14 | N.S. | N.S. | 0.074 ± 0.002 | 1.98 ± 0.03 | 61 ± 4 | 32 ± 2 |
| Igl | 2.2 ± 0.1 | 29 ± 5 | 78 ± 12 | N.S. | N.S. | 0.313 ± 0.007 | 2.3 ± 0.1 | 240 ± 40 | 10 ± 1 |
| 2Nal | 3.7 ± 0.3 | 7,100 ± 1,000 | 0.510 ± 0.04 | 5.3 ± 1.3 | 15,200 ± 6,500 | 0.35 ± 0.07 | 2.6 ± 0.2 | 1,800 ± 400 | 1.5 ± 0.2 |
| Bip | N.S. | N.S. | 0.0314 ± 0.0004 | N.S. | N.S. | 0.0178 ± 0.0004 | 2.53 ± 0.09 | 6,200 ± 800 | 0.41 ± 0.04 |
| Inp | 5.2 ± 0.4 | 65,000 ± 800 | 0.079 ± 0.004 | N.S. | N.S. | 0.03 ± 0.001 | 1.05 ± 0.06 | 5,600 ± 900 | 0.19 ± 0.02 |
| Amc | 5.7 ± 0.3 | 75,000 ± 7,000 | 0.076 ± 0.003 | 5.8 ± 0.3 | 27,000 ± 2500 | 0.213 ± 0.009 | 0.97 ± 0.04 | 1000 ± 200 | 0.94 ± 0.17 |
| 3Abz | 3.4 ± 0.4 | 3,100 ± 600 | 1.1 ± 0.1 | 4.8 ± 0.3 | 3000 ± 500 | 1.6 ± 0.2 | 1.08 ± 0.07 | 55 ± 12 | 20 ± 4 |
| Amb | N/A | 3,200 ± 1000 | N/A | N.S. | N.S. | 0.140 ± 0.007 | 1.6 ± 0.1 | 2100 ± 300 | 0.74 ± 0.06 |

N.S. No saturation, only $k_{\text{inact}}/K_i$ could be obtained. N/A, no time dependent inactivation was observed, data consistent with reversible inhibition.

Overall, changes in P2 do not have a big influence in *k*_inact_ with the exceptions of Asn, Phe(Me), and Tyr(NO_2_), which significantly increase *k*_inact_. Intriguingly, substrates containing the latter two P2 residues were the only substrates with a P2 aromatic residue that could be cleaved by DPAP3 with *k*_cat_/*K*_m_ > 2,000 M^−1^s^−1^ (Table 1). In terms of *K*_i_, we observed some SAR similarities between substrates and inhibitors: DPAP3 does not bind inhibitors with an N-terminal basic residue (Arg or Lys), but it is strongly inhibited (*K*_i_ < 35 nM) by aliphatic residues (Leu, nLeu, hAla, or Val). However, we measured potent inhibition of DPAP3 with aromatic residues in P2 such as Phe, Tyr or Trp (*k*_inact_/*K*_i_ ≥ 60,000 M^−1^s^−1^). Substrates with these P2 residues show relatively poor substrate turnover (*k*_cat_/*K*_m_ ≤ 2,000 M^−1^s^−1^). We also observed other discrepancies between substrates and inhibitors. For example, Thr in P2 results in a poor substrate (Fig. 1B) but a relatively potent inhibitor (*k*_inact_/*K*_i_ = 55,000 M^−1^s^−1^). Inversely, DPAP3 cleaves Phg-hPhe-ACC and Phe(Me)-hPhe-ACC with a *k*_cat_/*K*_m_ of 11,000 and 12,000 M^−1^s^−1^, respectively, but the *k*_inact_/*K*_i_ for the respective inhibitors are only 36 and 580 M^−1^s^−1^. As shown in Fig. 4A, there is not a clear correlation between *k*_cat_/*K*_m_ and *k*_inact_/K¡ for DPAP3 as a function of the P2 residue, however, P2 residues that make optimal substrates also make good inhibitors (Val, nVal, nLeu).

**Fig. 4.**
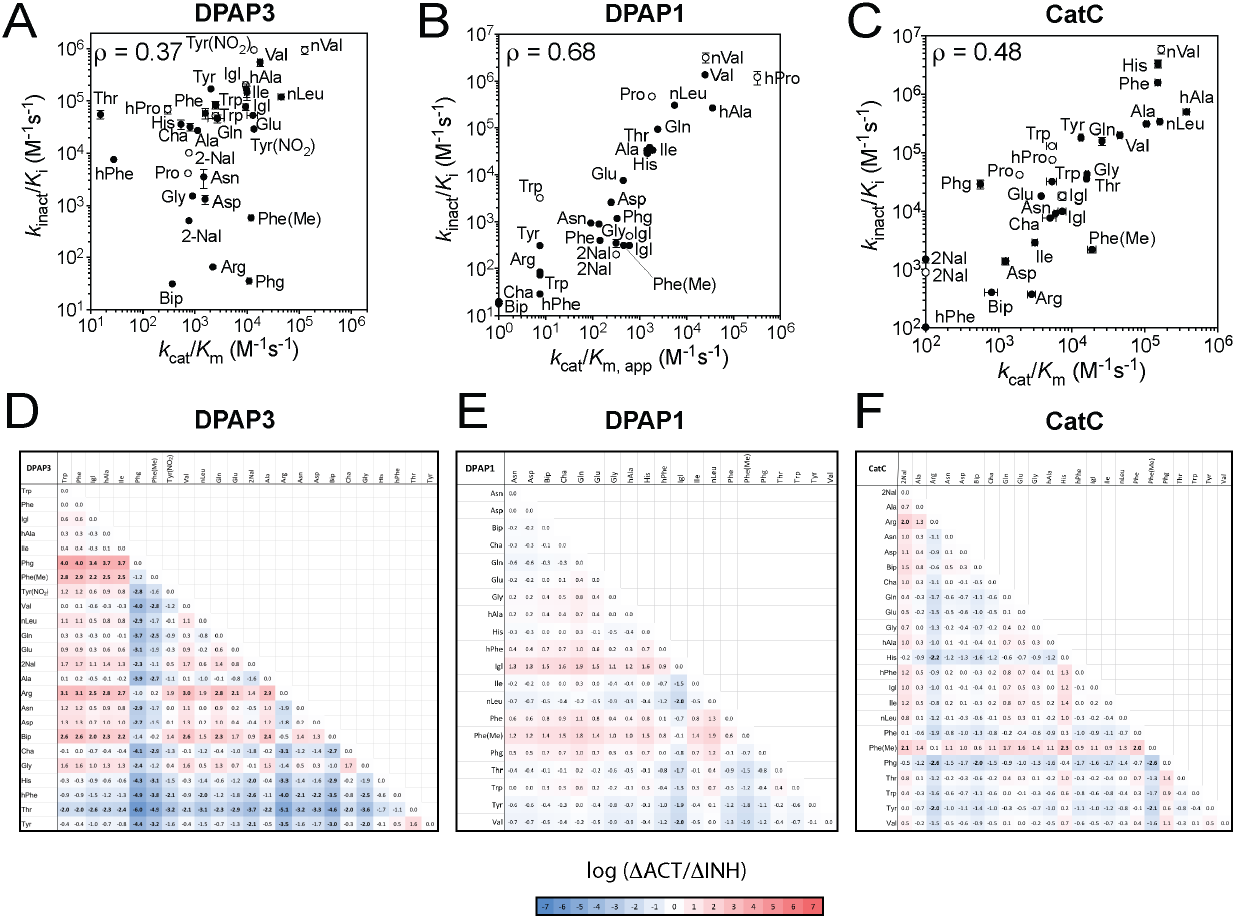
Comparison of catalytic efficiency and second order inhibition constants. (**A-C**) Comparison of *k*_inact_/*K*_i_ and *k*_cat_/*K*_m_ for DPAP3 (**A**), DPAP1 (**B**) and CatC (**C**) between vinyl sulfone inhibitors and P2 substrate library. For DPAP1, apparent *k*_cat_/*K*_m_(*k*_cat_/*K*_m,app_) were calculated based on previously reported turnover rates at 1 μM[34]. *k*_cat_/*K*_m,app_ were calculated similarly for DPAP3 and CatC for P2 substrates whose activity was too low to obtain accurate Michaelis Menten parameters (i.e., substrates not present in Tables 1 and S2). Filled circles correspond to compounds belonging to the vinyl sulfone library (compounds in Table 2), and empty circles to inhibitors having a phenyl group in P1’ (Table 3). The P2 residue is labelled next to each data point. Pearson correlation coefficients (*ρ*) are shown for each protease. (**D-F**) Comparison of changes in substrate turnover relative to inhibitors potency for any pair of P2 residues. The log value of ΔACT/ΔINH (Eq. 2) calculated for DPAP3 (**D**), DPAP1 (**E**) and CatC (**F**) are shown has a heat map with values above and below zero in red and blue, respectively. Each pairwise value showing more than a 100-fold discrepancy between activity and inhibition (ΔACT/ΔINH > 100 or < 0.01) is highlighted in bold.

### Correlation between substrate turnover and inhibition for DPAP1 and CatC

To determine whether the lack of correlation between *k*_cat_/*K*_m_ and *k*_inact_/*K*_i_ observed for DPAP3 is a common feature in DPAPs, we calculated apparent *k*_cat_/*K*_m_ values for DPAP1 for the P2 substrate library based on the activity measurements previously reported at 1 μM and the *k*_cat_/*K*_m_ for hPro-hPhe-ACC[39]. We also measured accurate Michaelis Menten parameters for CatC for the P2 substrate library (Table S1 and Fig. S3), and *k*_inact_ and *K*_i_ values for DPAP1 and CatC for the P2 vinyl sulfone library (Table 2 and Fig. S2). As shown in Fig. 4B, we observed a good correlation between substrate turnover and *k*_inact_/*K*_i_ for DPAP1, but for CatC we observed some discrepancies (Fig. 4C), albeit not as pronounced as in the case of DPAP3. For example, the CatC *k*_cat_/*K*_m_ for Phe(Me)-hPhe-ACC is 30-fold higher than for Phg-hPhe-ACC while the *k*_inact_/*K*_i_ for Phe(Me)-hPhe-VS is 13-fold lower than for Phg-hPhe-VS, thus resulting in a 400-fold discrepancy in the changes in *k*_cat_/*K*_m_ and *k*_inact_/*K*_i_ values between these two P2 residues.

A systematic way to visualize discrepancies between substrate turnover and inhibition is to compare the fold different in *k*_cat_/*K*_m_ with that in *k*_inact_/*K*_i_ for any two P2 residues (Eq. 2).

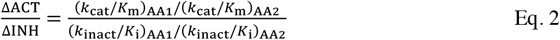

We performed these pairwise calculations for each of the DPAPs studied here and have presented the results as heat maps in Fig. 4D-F. We observed significant and numerous discrepancies between substrates and inhibitors for DPAP3 (30 % of pairwise ΔACT/ΔINH > 100 or < 0.01), almost no discrepancies for DPAP1 (only 1% ÁACT/AINH > 10^2^ or < 10^−2^), and only a few for CatC (4% of ΔACT/ΔINH > 100 or < 0.01). Overall, this study indicates that the level of correlation between *k*_cat_/*K*_m_ and *k*_inact_/*K*_i_ is protease dependent.

### Development of DPAP1 and DPAP3-selective inhibitors

We next synthesized several inhibitors to determine whether the optimal nLeu(o-Bzl) P1 residue identified from the substrate screen could be used to increase the potency and specificity of inhibitors towards DPAP1 or DPAP3. We selected P2 AAs that were predicted to provide specificity towards DPAP1 (Pro and hPro) or DPAP3 (aromatic residues: Tyr(NO_2_), Trp, Igl and 2Nal) based on our substrate and inhibitor screening results. We also included in our analysis the previously synthesized compound JCP410 (nVal-hPhe-VS)[26], since nVal is one of the best P2 residues identified from the substrate screen. We determined the inhibition constants of these compounds for DPAP1, DPAP3, and CatC (Table 3 and Figs. 3 and S2). The major structural difference between these compounds and the inhibitor library is that they have a phenyl group in P’ instead of a long aliphatic linker (Fig. 3A). This change usually increases the potency of compounds except in the context of a P2 Trp for DPAP3 or P2 Tyr(NO_2_)/2Nal for CatC (Tables 2 and 3). These exceptions indicate some level of interdependence between the prime and non-prime sites of DPAPs.

**Table 3.** Inhibition constants for optimal vinyl sulfone inhibitors

| Inhibitor Name | DPAP3 |  |  | DPAP1 |  |  | CatC |  |  | Specificity Ratio<sup>a</sup> |  |  |
| --- | --- | --- | --- | --- | --- | --- | --- | --- | --- | --- | --- | --- |
| | $k_{\text{inact}} \cdot 10^{-3}$<br>(s<sup>-1</sup>) | $K_i$<br>(nM) | $k_{\text{inact}}/K_i \cdot 10^{-3}$<br>(M<sup>-1</sup>s<sup>-1</sup>) | $k_{\text{inact}} \cdot 10^{-3}$<br>(s<sup>-1</sup>) | $K_i$<br>(nM) | $k_{\text{inact}}/K_i \cdot 10^{-3}$<br>(M<sup>-1</sup>s<sup>-1</sup>) | $k_{\text{inact}} \cdot 10^{-3}$<br>(s<sup>-1</sup>) | $K_i$<br>(nM) | $k_{\text{inact}}/K_i \cdot 10^{-3}$<br>(M<sup>-1</sup>s<sup>-1</sup>) | DPAP1/<br>DPAP3 | DPAP1/<br>CatC | DPAP3/<br>CatC |
| Pro-hPhe-VS | 3.5 ± 0.2 | 860 ± 100 | 4.1 ± 0.3 | 2.1 ± 0.2 | 4.6 ± 0.7 | 470 ± 35 | 1.45 ± 0.07 | 34 ± 5 | 42 ± 5 | **115** | 11 | 0.10 |
| hPro-hPhe-VS | 2.1 ± 0.1 | 30 ± 7 | 69 ± 13 | 4.8 ± 0.6 | 3.9 ± 1.5 | 1,230 ± 400 | 1.67 ± 0.08 | 22 ± 3 | 76 ± 0.8 | 18 | 16 | 0.91 |
| hPro-nLeu(o-Bzl)-VS | 2.0 ± 0.1 | 39 ± 6 | 53 ± 6.5 | 2.9 ± 0.3 | 0.8 ± 0.2 | 3,560 ± 150 | 1.67 ± 0.03 | 11.2 ± 0.6 | 150 ± 6 | 68 | 24 | 0.35 |
| Tyr(NO<sub>2</sub>)-hPhe-VS | 2.26 ± 0.05 | 2.4 ± 0.2 | 950 ± 74 | 3.8 ± 1 | 25,000 ± 1,400 | 0.15 ± 0.05 | 3.2 ± 0.6 | 58,000 ± 20,000 | 0.054 ± 0.008 | **0.00016** | 2.8 | **17,600** |
| Tyr(NO<sub>2</sub>)-nLeu(o-Bzl)-VS | 2.7 ± 0.2 | 53 ± 10 | 51 ± 10 | 2.6 ± 0.3 | 2,400 ± 900 | 1.1 ± 0.3 | 2.9 ± 0.2 | 7,700 ± 1,400 | 0.38 ± 0.05 | 0.032 | 2.9 | 89 |
| 2Nal-hPhe-VS | 2.59 ± 0.07 | 254 ± 16 | 10.2 ± 0.5 | 6.7 ± 0.2 | 33,000 ± 2,000 | 0.2 ± 0.005 | 3.7 ± 0.2 | 4,000 ± 700 | 0.9 ± 0.1 | 0.02 | 0.22 | 11 |
| 2Nal-nLeu(o-Bzl)-VS | 6.0 ± 0.7 | 2,800 ± 600 | 2.2 ± 0.3 | 5.0 ± 1.7 | 27,000 ± 16,000 | 0.19 ± 0.05 | 2.4 ± 0.6 | 1,740 ± 140 | 1.37 ± 0.09 | 0.14 | 0.14 | 1.0 |
| Igl-hPhe-VS | 2.1 ± 0.7 | 10 ± 1 | 205 ± 22 | 2.7 ± 0.1 | 5,400 ± 500 | 0.50 ± 0.03 | 5.8 ± 0.3 | 320 ± 70 | 18 ± 3 | **0.0024** | 0.028 | 11.4 |
| Igl-nLeu(o-Bzl)-VS | 5.6 ± 0.4 | 160 ± 30 | 35 ± 4.5 | 4.1 ± 0.2 | 4,800 ± 500 | 0.86 ± 0.05 | 1.48 ± 0.06 | 11 ± 2 | 135 ± 14 | 0.037 | **0.0064** | 0.17 |
| L-Trp-hPG-VS | 3 ± 1 | 330 ± 150 | 9.1 ± 1.5 | 2.1 ± 0.2 | 2,800 ± 450 | 0.74 ± 0.07 | 1.73 ± 0.03 | 28 ± 2 | 61 ± 4 | 0.12 | 0.012 | 0.10 |
| L-Trp-hPhe-VS | 3.3 ± 0.4 | 65 ± 15 | 52 ± 7 | 1.7 ± 0.1 | 540 ± 80 | 3.20 ± 0.3 | 2.2 ± 0.09 | 16 ± 3 | 131 ± 15 | 0.094 | 0.024 | 0.26 |
| L-Trp-nLeu(o-Bzl)-VS | 3.2 ± 0.3 | 4 ± 1 | 826 ± 193 | 2.2 ± 0.1 | 380 ± 60 | 5.8 ± 0.6 | 1.74 ± 0.03 | 4.9 ± 0.3 | 355 ± 18 | **0.0070** | 0.016 | 2.3 |
| D-Trp-hPhe-VS | 1.7 ± 0.6 | 5,500 ± 3,000 | 0.32 ± 0.08 | 2.7 ± 0.2 | 105,000 ± 10,000 | 0.026 ± 0.001 | 2.6 ± 0.1 | 4,200 ± 600 | 0.62 ± 0.07 | 0.12 | 0.042 | 0.34 |
| D-Trp-nLeu(o-Bzl)-VS | 3.6 ± 0.2 | 7,600 ± 600 | 0.47 ± 0.01 | 5.5 ± 0.7 | 127,000 ± 20,000 | 0.044 ± 0.002 | 1.97 ± 0.06 | 1,100 ± 100 | 1.7 ± 0.1 | 0.094 | 0.026 | 0.28 |
| nVal-hPhe-VS | 2.6 ± 0.4 | 2.8 ± 0.9 | 940 ± 160 | 2.0 ± 0.1 | 0.6 ± 0.2 | 3,160 ± 750 | 2.2 ± 0.1 | 0.38 ± 0.08 | 5,830 ± 970 | 3.36 | 0.54 | 0.16 |

<sup>a</sup>Specificity ratio was calculated based on $k_{\text{inact}}/K_i$ values; numbers highlighted in bold indicate more than 100-fold selectivity for the indicated enzymes.

Compared to hPhe, P1 nLeu(o-Bzl) decreases *k*_inact_/*K*_i_ value for DPAP3 by 4 to 18-fold except in the context of a P2 Trp where it increases it by 24-fold, or a P2 hPro where there is no significant change (Table 3). These differences are mainly due to changes in *K*_i_ rather than *k*_inact_ and likely reflect cooperativity between the S1 and S2 pockets of DPAP3. However, in the case of DPAP1 and CatC, replacement of P1 hPhe with nLeu(o-Bzl) decreases *K*_i_. Because of this P1-P2 interdependence, the most potent inhibitors for DPAP3 are either a combination of P2 Tyr(NO_2_) and P1 hPhe, i.e. SAK1, or P2 Trp and P1 nLeu(o-Bzl), resulting in *k*_inact_/*K*_i_ values close to 10^6^ M^−1^s^−1^. Importantly, nVal-hPhe-VS is as potent as these two inhibitors (*k*_inact_/*K*_i_ = 940,000 M^−1^s^−1^), confirming that optimal inhibitors can be designed based on the structure of optimal substrates. These high second order rate constants are mainly driven by low *K*_i_ values (< 5 nM). While Trp-nLeu(o-Bzl), Igl-hPhe-VS and Tyr(NO_2_)-hPhe-VS show more than 100-fold selectivity for DPAP3 over DPAP1, only the latter is selective for DPAP3 compared to CatC (Table 3). On the other hand, nVal-hPhe-VS is equally potent for all DPAPs (*k*_inact_/*K*_i_ = 1 10^6^, 3.2·10^6^ and 5.8·10^6^ M^−1^s^−1^ for DPAP3, DPAP1 and CatC, respectively) making it a highly potent but non-selective pan-DPAP inhibitor Pro-hPhe-VS (SAK2) is 100-fold more selective towards DPAP1 than DPAP3, but only shows a 10-fold selectivity for DPAP1 compared to CatC. While replacing the P2 Pro of SAK2 with hPro increases the potency of the inhibitors towards DPAP1, this also results in some loss of specificity (Table 3). Overall, we were able to increase the potency of SAK2 (Pro-hPhe-ACC) towards DPAP1 by 7-fold by using the optimal P1 (nLeu(o-Bzl)) and P2 (hPro) residues identified from the substrate library screen[34], making hPro-nLeu(o-Bzl)-VS the most potent DPAP1 inhibitor identified so far (*k*_inact_/*K*_i_ = 3.610^6^ M^−1^s^−1^)[25,26,42]. However, this 7-fold increase in inhibitor potency between Pro-hPhe-VS and hPro-nLeu(o-Bzl)-VS was much lower than expected based on the 20- and 30-fold increase in susbtrate turnover reported between P1 hPhe and P1 nLeu(o-Bzl), and between P2 Pro and P2 hPro, respectively[34].

### Testing inhibitors specificity in live parasites

We then tested the potency and selectivity of our DPAP1 and DPAP3 specific inhibitors in live parasites using the FY01 activity-based probe (ABP) in a competition labelling assay. ABPs are small molecule reporters of activity that use the catalytic mechanism of the targeted enzyme to covalently modify its active site. A reporter tag, usually a fluorophore or a biotin, allows visualization and quantification of the labelled enzyme in a gel-based assay[43]. FY01 is a cell-permeable fluorescent ABP that was initially developed for CatC[44] but it also labels *Plasmodium* DPAPs and the falcipains in live parasites[25,26]. The falcipains (FPs) are clan CA Cys proteases involved in haemoglobin degradation (FP2, FP2’ an FP3)[45] and possibly RBC invasion (FP1)[46,47]. Binding of inhibitors into the active site of any of these Cys proteases prevents probe labelling resulting in the disappearance of a fluorescent band a SDS-PAGE gel.

Live parasites were treated with different concentrations of inhibitor for 30 min, and the residual level of DPAPs and FPs activities labelled with FY01 (Fig. S4) and quantified by densitometry. Dose response curves are shown in Fig. 5 and IC_50_ values reported in Table 4. Inhibitors with a P2 Pro or hPro are equally potent and inhibit DPAP1 at mid nanomolar concentrations. However, the P2 Pro makes the inhibitor more selective by blocking inhibition of the FPs. Compounds with a P2 Trp or Tyr(NO_2_) are highly specific for DPAP3 in intact parasites. Surprisingly, Trp-hPG-VS is by far the most potent DPAP3 inhibitor in intact parasites (IC_50_ = 1.4 nM) despite being 5 to 100-fold less potent than Trp-nLeu(o-Bzl)-VS or Trp-hPhe-VS against DPAP3 (see *k*_inact_/*K*_i_ values in Table 3). This suggests that either this compound is metabolically more stable and/or that decreasing the hydrophobicity of the P1 residue enhances the cell permeability of the compound. Indeed, compounds need to cross four membranes to reach DPAP3: the RBC, parasitophorous vacuole, and parasite plasma membranes, plus the membrane of the apical organelle where DPAP3 resides[21]. (The parasitophorous vacuole is a membrane bound structure within which the parasite growth and multiply isolated from the RBC cytosol.) It is also likely that the apparent increased potency of Trp-hPG-VS in live parasites might be due to its accumulation into the DPAP3 acidic organelle via protonation of its free amine. However, we predict that this lysosomotropic effect likely occurs for all DPAP inhibitors presented here.

**Fig. 5.**
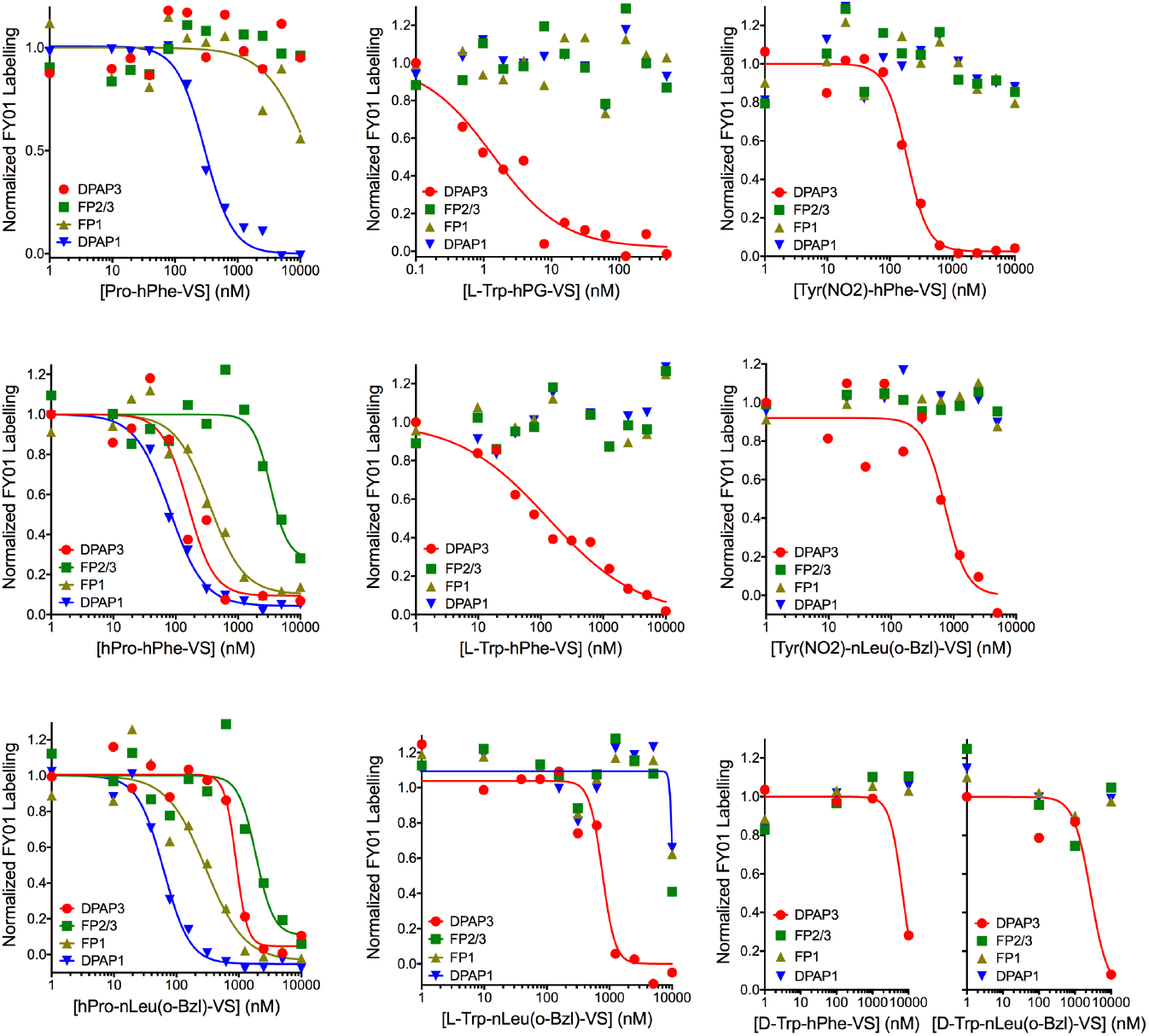
Selective inhibition of DPAP1 or DPAP3 in live parasites. Intact mature schizonts were pre-treated for 30 min with a dose response of inhibitor followed by 1 h treatment with FY01 to label the residual level of DPAP1, DPAP3, FP1 and FP2/3 activities. Fluorescents bands corresponding to the different cysteine proteases labelled by the probe (Fig. S4) were quantifies by densitometry using ImageJ, and the fluorescence values normalized to the DMSO control. IC_50_ values are reported in Table 4. Note that control inhibitors containing *D*-Trp are more than 1000-fold less potent than those with *L*-Trp.

**Table 4.** Cysteine proteases IC_50_ values in live parasites.

| Inhibitor | IC<sub>50,DPAP3</sub> (nM) | IC<sub>50,DPAP1</sub> (nM) | IC<sub>50,FP1</sub> (nM) | IC<sub>50,FP2/3</sub> (nM) |
| --- | --- | --- | --- | --- |
| hPro-hPhe-VS | 160 ± 50 | 80 ± 7 | 350 ± 80 | 3,300 ± 1,000 |
| Pro-hPhe-VS | > 10,000 | 275 ± 25 | ~ 10,000 | > 10,000 |
| hPro-nLeu(o-Bzl)-VS | 900 ± 100 | 62 ± 7 | 300 ± 100 | 1,900 ± 500 |
| Tyr(NO<sub>2</sub>)-hPhe-VS | 190 ± 20 | > 10,000 | > 10,000 | > 10,000 |
| Tyr(NO<sub>2</sub>)-nLeu(o-Bzl)-VS | 730 ± 170 | > 10,000 | > 10,000 | > 10,000 |
| L-Trp-hPG-VS | 1.4 ± 0.4 | > 10,000 | > 10,000 | > 10,000 |
| L-Trp-hPhe-VS | 130 ± 60 | > 10,000 | > 10,000 | > 10,000 |
| L-Trp-nLeu(o-Bzl)-VS | 760 ± 150 | ~ 10,000 | ~ 10,000 | ~ 10,000 |
| D-Trp-hPhe-VS | 6,800 ± 1,600 | > 10,000 | > 10,000 | > 10,000 |
| D-Trp-nLeu(o-Bzl)-VS | 2,700 ± 1,600 | > 10,000 | > 10,000 | > 10,000 |

### Correlation between PS-SCL of substrates and inhibitors for other Cys proteases

To determine whether the discrepancy observed between substrate and inhibitor specificities for DPAP3 (Fig. 4) is a phenomenon observed in other proteases, we look at previously published literature about two of the most commonly studies families of Cys proteases, i.e. caspases and cathepsins. Several studies on caspases[48,49] and cathepsins[50] have shown that inhibitors designed based on the structure of specific substrates do not retain their selectivity. Also, the sequence of an optimal substrate or inhibitor might render the equivalent inhibitor or substrate completely inactive[49–51]. We therefore compared previously published specificity data obtained from PS-SCL libraries of ACC substrates and covalent inhibitors for these two protease families (Fig. 6).

**Fig. 6.**
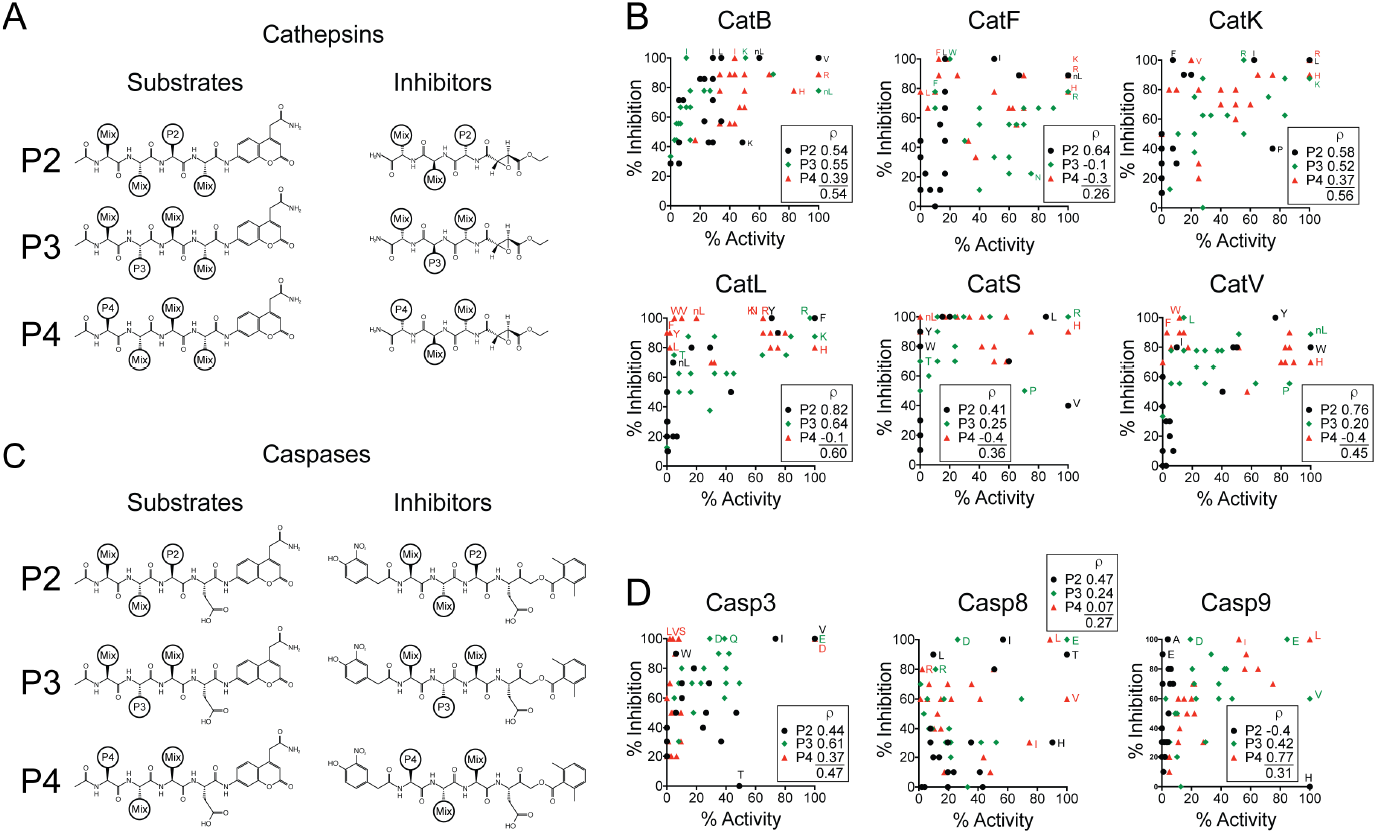
Comparison of substrate and inhibitor specificity for cathepsins and caspases. (**A** and **C**) Structure of the PS-SCL of ACC substrates and inhibitors used to determine the specificity of cathepsins and caspases. For the cathepsins, substrates were screened at 250 μM, and the inhibitors at 10 μM using an ABP competition assay. For caspases, substrates were screened at 50 μM and inhibitors at 50-500 nM. (**B** and **D**) Correlation between substrate turnover and inhibition. For each protease and P2-P4 position, the maximum level of activity and inhibition at each AA position was normalized to 100 %. Pearson correlation coefficients (*ρ*) are shown next to the legend of each graph for each position. The lower *ρ* value was obtained by combining all the P2-P4 data points for each protease. Residues showing maximum activity or inhibition are labelled at each position along with specific AAs that illustrate particularly bad correlation between the levels of activity and inhibition.

To simplify our analysis, we only compared data obtained from natural amino acids. For Cys cathepsins (CatB, CatF, CatK, CatL, CatS, and CatV), we compared the substrate specificity data obtained from the Craik lab[50] with the inhibitor specificity data obtained from the Bogyo lab using an epoxysuccinate library[36] (Fig. 6A-B). For caspases, we compared the specificity obtained for caspases 3, 8 and 9 (Casp3, Casp8 and Casp9, respectively) using a library of acyloxymethyl ketone inhibitors from the Bogyo lab[51], with the substrate specificity results from the Drag lab [52] (Fig. 6C-D). For each protease, we normalized the level of activity and inhibition to that of the best natural amino acid at each P2-P4 position. Although it is difficult to draw specific conclusions from these PS-SCL results because each activity or inhibition value represent the effect of a pool of 400-8000 different molecules rather than an individual one, the lack of correlation between the levels of activity and inhibition shown in Fig. 6 is nonetheless quite surprising. Indeed, for most proteases and P2-P4 positions, the Pearson coefficients are below 0.7. Similar to what we observed for DPAPs (Fig. 4), the peptide sequence of inhibitors that is equivalent to that of optimal substrates generally yields very potent inhibitors. This is true for most proteases and for each position (with the exception of P2 in Casp9). However, for all proteases, 90-100 % inhibition was achieved with peptide sequences that were poorly cleaved as substrates (< 10 % activity). Also, at each P2-P4 position, we could observe good inhibition with sequences belonging to poor substrates. Interestingly, for most proteases only a few peptide sequences showed good turnover rates but poor inhibition, with the exception of Casp8, and the P3 position for CatF. There are also a few striking individual residues, such as P2 His for caspases 8 and 9, or P2 Thr for Casp3 and P2 Val for CatS, that show good activity but no inhibition. Finally, we would also like to draw attention to the fact that the P2 and P4 positions are particularly important for modulating the potency of inhibitors against Casp9 and Casp3, respectively. However, these positions have little effect on substrate turnover.

That said, there are substantial structural differences between the substrate and inhibitor libraries shown in Fig. 6 that can explain this lack of correlation. For epoxysuccinate inhibitors, the polarity of the peptide backbone goes in the opposite direction as that of natural substrates, these compounds do not have a P1 residue, and the warhead strands the S1 and S1’ pocket. This can have very drastic influence on the preferred amino acid residue at each site, especially at the P2 and P3 position. Also, the position of the site of nucleophilic attack by the catalytic Cys at one of the epoxide carbons does not match the position of the scissile bond carbonyl in a peptide substrate and can therefore explain the differences in specificity between substrate and inhibitors. This is therefore an excellent example showing how the choice of warhead can have drastic consequences when the structural information of a substrate wants to be used to develop covalent inhibitors.

As pointed above, differences in specificity between substrate and inhibitors have been pointed out previously for caspases. A possible explanation to explain the differences observed in Fig. 6 is that the P4 capping group is different between the substrate (acetyl group) and inhibitor (2-(4-hydroxy-3-nitrophenyl)-acetyl group) libraries. This might strongly influence how side chains, specially at the P4 position, bind into the different active site pockets. Indeed, using different capping group in PS-CSL of substrates has been shown to result in different amino acid preferences in the S4 pocket of the Zika virus NS2B-NS3 protease[53,54]. Also, if cooperativity exist between the P4 and other positions, the influence of the capping group on the S4 pocket specificity will also influence that of other pockets. Finally, the difference in specificity might be due to cooperativity between the prime and non-prime sites. In this case the choice of electrophile used might change the amino acid preference.

## Discussion

This study provides the first characterization of the specificity of DPAP3, a cysteine protease important for efficient invasion of RBCs by the malaria parasite[21]. DPAP3 is a highly efficient proteolytic enzyme showing similar *k*_cat_ and *k*_cat_/*K*_m_ values than either DPAP1 or CatC when optimal substrates are used (Table 1). In general, DPAP3 has a narrower substrate specificity than either DPAP1 or CatC, and it preferentially cleaves substrates with aliphatic residues at the N-terminus. Our study also shows similar P1 substrate specificity across all DPAPs, i.e. a strong preference for basic or long aromatic residues. DPAP3 was the only DPAP able to cleave some substrates with aromatic P2 residues (Tyr, Phe(Me), Phg), albeit with relatively low turnover rates. Surprisingly, vinyl sulfone inhibitors with aromatic residues are as potent as compounds with optimal P2 residues identified from the substrate screen, but in addition, these compounds are highly specificity for DPAP3 compared to DPAP1, CatC, or other malarial cysteine proteases (FPs). The previously described DPAP1 inhibitor SAK2 (Pro-hPhe-VS) shows the greatest specificity for DPAP1 in live parasites. Incorporation of optimal P1 (nLeu(o-Bzl)) and P2 (hPro) residues identified from the substrate screens improves the potency of DPAP1 inhibitors both *in vitro* and in live parasites, but also results in some loss in specificity (Table 3 and Fig. 5).

Despite being highly potent and specific DPAP1 and DPAP3 inhibitors, these compounds only show antiparasitic activity at mid to high micromolar concentrations[21,25,26], probably due to metabolic stability issues. Indeed, we have previously shown that this is the case for Pro-hPhe-VS, which is not able to sustain target inhibition in live parasites[25]. A possible cause for this instability is the presence of multiple aminopeptidases in the malaria parasite that might cleave the amide bond of these compounds[55], thus preventing them to bind into the DPAPs active sites. Nonetheless, this study provides a very strong SAR foundation to develop potent non-peptidic inhibitors able to sustain DPAPs inhibition. From a drug development point of view, our SAR studies indicate that inhibitors with short aliphatic P2 residues strongly inhibit both DPAPs (Fig. 4E), indicating that potent pan-DPAP inhibitors can easily be developed. Unfortunately, we did not identify any clear P1 or P2 residue that would discriminate malarial DPAPs from host CatC. Therefore, further studies are required to determine whether differences in specificity in the prime binding pockets can be exploited to develop parasite specific inhibitors.

It is however important to point out that given the short-term treatment requirement for antimalarial therapy (single dose or less than 3 doses in 3 days), we think it is unlikely that inhibition of CatC would lead to adverse side effects. Firstly, highly specific DPAP inhibitors might not be necessary given that a high level (> 95 %) of sustained CatC inhibition is required to see a decrease in the activity of serine proteases activated by CatC[56]. Secondly, activation of granule serine proteases by cathepsin C takes place during cell differentiation in the bone marrow, and a decrease in the level of serine proteases activity in circulating immune cells is only achieved after more than 2 weeks of daily treatment with CatC inhibitors[19]. And thirdly, no signs of toxicity were observed in phase I clinical trials when volunteers were treated daily for more than 3 weeks with CatC inhibitors, albeit some on-target side effects such as plantar and palmar epithelial desquamation were observed in some instances[18,19]. However, these side effects were not observed in volunteers that received a single dose of CatC inhibitor, nor within the first week of daily treatments.

Positional scanning substrate libraries have been used over the last 20 years[29–31] to determine the specificity of proteases and guide the synthesis of inhibitors. Here we have shown that vinyl sulfone inhibitors containing P1 and P2 residues corresponding to optimal DPAP1 or DPAP3 substrates result in extremely potent inhibitors. However, our studies clearly identified differences in specificity between substrates and inhibitors, especially for DPAP3, but also for CatC and other cysteine proteases (Figs. 4 and 6). A reason why this phenomenon might not have been more broadly reported in the literature is because, in general, either substrate or inhibitor libraries are used to determine the specificity of a protease, but not both. Also, inhibitors are not usually designed based on the structure of non-optimal substrates, and as pointed above, inhibitors that mimic optimal substrates are generally very potent. That said, there are multiple reasons that can account for discrepancies in specificity between substrates and inhibitors:

First, although the substrate and inhibitor libraries used in this study have equivalent P1 and P2 residues, the structural features that bind into the S’ pockets are quite different. Therefore, if the specificity of a protease shows interdependence between its prime and nonprime binding pockets, it might explain differences in specificity. This is likely the case for DPAPs since we observed a 50-fold increase in *k*_cat_ between the Phe-Arg-ACC and Phe-Arg-βNA substrates, which only differ in the structure of the fluorophore that binds in the S1’ pocket (Table 1). Also, while we observed very good correlation between *k*_cat_/*K*_m_ and *k*_inact_/*K*_i_ of DPAP1 for the P2 substrate and VS library (compounds in Table 2), in the context of a phenyl group in P1’ (inhibitors in Table 3), we observed clear discrepancies between substrates and inhibitors (Fig. 4B).

Second, the position of the electrophilic warhead within the active site might differ from that of the scissile bond in a substrate, especially in terms of distance and orientation relative to the catalytic Cys. This positioning might be differently affected by changes in the P1 and P2 residues of substrates and inhibitors. Indeed, a recent study on caspases has shown that acyloxymethyl ketone covalent inhibitors might act through a reversible mechanism even if they are designed based on the sequence of optimal substrates[49]. Also, ABPs designed to profile deubiquitinating proteases by conjugating an electrophile to the C-terminal of ubiquitin have been shown to label different subset of enzymes depending on the warhead used[57].

Third, substrate turnover by Cys and Ser proteases requires two different chemical steps: First, nucleophilic attack of the peptide bond by the catalytic residue to form the acyl intermediate and release of the C-terminal product of proteolysis; and second, hydrolysis of the acyl intermediate by an activated water molecule to reconstitute the free enzyme and release the N-terminal product of the reaction (Scheme 1). Therefore, peptide sequences that are poorly turned over because of a very slow acyl intermediate hydrolysis step might still be very good to design covalent inhibitors. For example, a substrate that displaces the catalytic water from the active site upon formation of the acyl intermediate will be very poorly turned over.

**Scheme 1.**
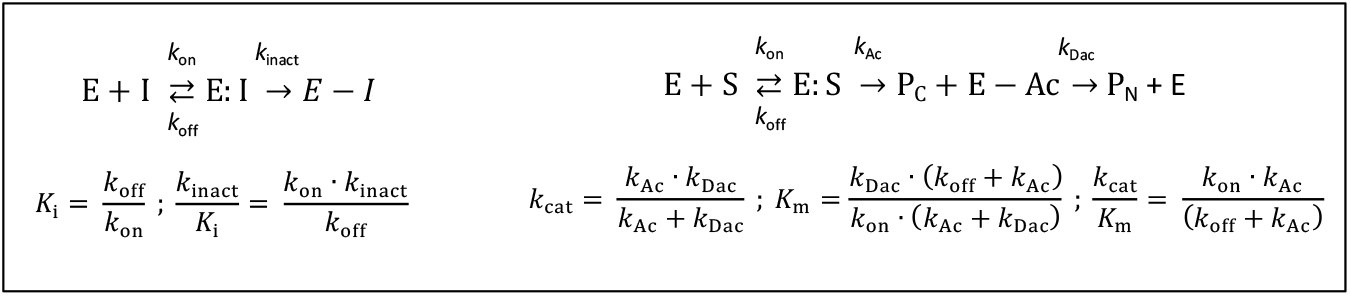
Comparison between irreversible inhibition and substrate turnover kinetic constants. E, I, E:I and E-I represent free enzyme, inhibitor, inhibitor associated with the enzyme, and enzyme covalently modified by the inhibitor, respectively. S, E:S, E-Ac, P_C_ and P_N_ represent the substrate, the Michaelis Menten enzyme-substrate complex, the acyl intermediate, and the C- and N-terminal products of proteolysis, respectively; *k*_on_ and *k*_off_ the association and dissociation rate constant for substrates or inhibitors; and *k*_Ac_ and *k*_Dac_ the kinetic constants for formation of the acyl intermediate and its hydrolysis, respectively.

And fourth, the reaction mechanism between covalent inhibition and substrate turnover are quite different making *k_cat_, *K*_m_, and *k*_cat_/*K*_m_ not directly comparable with *k*_inact_, *K*_i_*, and *k*_inact_/*K*_i_ (Scheme 1). *k*_cat_, *K*_m_, and *k*_cat_/*K*_m_ are empirical parameters obtained under steady state conditions while *k*_inact_, *K*_i_, and *k*_inact_/*K*_i_ are real kinetic and thermodynamic constants that cannot be measured under steady state conditions since covalent inhibitors deplete the concentration of free enzyme over time. *k_cat_* might be equivalent to *k*_inact_ only if *k*_Ac_ ≪ *k*_Dac_, i.e. formation of the acyl intermediate is the rate limiting step; *K*_m_ might be equivalent to *K*_i_ only if *k*_Ac_ ≪ *k*_off_ and *k*_Ac_ ≪ *k*_Dac_; and *k*_cat_/*K*_m_ might be equivalent to *k*_inact_/*K*_i_ only if *k*_off_ ≫ *k*_Ac_. Although these conditions might be met for certain substrates, the relative magnitudes of all these rate constants depend on the substrate sequence. For example, detailed pre-steady state kinetics and kinetic isotope effects studies on CatC have shown that the rate limiting step in the turnover of dipeptidic AMC substrates can be either formation of the acyl intermediate, its hydrolysis, or a contribution of both, depending on the nature of the P1 residue[58,59].

Although several studies have compared the potency of inhibitors to the turnover of equivalent substrates for selected peptide sequences, to the best of our knowledge this is the first study that systematically compares the potency of a peptide-based covalent inhibitor library to the turnover efficiency of an equivalent substrate library. We think that the discrepancies observed here between *k*_cat_/*K*_m_ and *k*_inact_/*K*_i_ are likely to be present in other proteases, as shown for cysteine cathepsins and caspases, but the level of discrepancy will be dependent on the protease studied as well as on the type of substrate and covalent inhibitor that are being compared, especially if a there is a high level of cooperativity between the prime and non-prime binding pockets.

Overall, our detailed study on DPAPs specificity and our analysis of specificity results obtained from PS-SCL clearly indicate that there can be very significant differences in specificity between substrates and covalent inhibitors. Although it is now well established that highly potent inhibitors can be developed based on the structure of optimal substrates, this might sometimes result in some loss of specificity. This study clearly demonstrates that optimal inhibitors with improved specificity can be developed based on the structure of relatively poor substrates. Therefore, we strongly recommend using PS-SCL libraries of inhibitors rather than substrates if the goal is to design specific inhibitors and ABPs, or select warheads and capping groups (N- and C-terminal) that mimic as well as possible the structure of substrates used to determine the specificity of a protease to design covalent inhibitors since these will be less likely to influence inhibitor specificity.

## Materials and Methods

Reagents. The syntheses of the DPAP substrate library and that of Val-Arg-ACC, Phe-Arg-ACC, Tyr(NO_2_)-hPhe-ACC[39] and (PR)_2_Rho[40] have been previously described. The syntheses of additional substrates used in this study are described in the supplementary methods and were synthesized following previously published methods[34],[60]. The syntheses of the vinyl sulfone inhibitor library, SAK2, SAK1, *L*-WSAK and *D*-WSAK were also previously described[21],[26]. The additional DPAP inhibitors used in this study were synthesized following similar methods[61]. The synthesis and characterization of these inhibitors and their synthetic intermediates are described in detail in the supplementary methods. The Phe-Arg-βNA and Gly-Arg-AMC substrates were purchased from Sigma. Recombinant DPAP3 was expressed in insect cells using the bacculovirus system as recently described[21]. Bovine CatC was purified to homogeneity from spleen by modification of a method described previously[62],[63].

### Recombinant DPAP3 active site titration

Recombinant DPAP3 was expressed in SF9 insect cells and purified from the culture supernatant by sequential ion exchange, Ni-NTA, and size exclusion chromatography as previously described[21]. To accurately determine the concentration of active DPAP3 in our enzyme stocks we used the FY01 ABP. Because ABPs only react with the active form of an enzyme and covalently modify its active site, in this case the catalytic Cys, they can be used to perform accurate active site titrations.

Our stock of DPAP3 was diluted 20-fold in assay buffer (100 mM sodium acetate, 100 mM NaCl, 5 mM MgCl_2_, 1 mM EDTA, 0.1 % CHAPS, and 5 mM DTT at pH 6), pretreated for 30 min with DMSO or 1 μM SAK1 (Tyr(NO_2_)-hPhe-VS), and DPAP3 labelled with 1 μM FY01 for 1 h at RT. These samples were run on a SDS-PAGE gel along with a serial dilution of free probe (1.5-100 nM). In-gel fluorescence was measured using a Bio-Rad PharosFX flat-bed scanner and the intensity of the fluorescent bands quantified using ImageJ. The fluorescent signal from the free probe was used as a calibration curve and compared to the difference in signal between the DMSO and SAK1 treated DPAP3 (Fig. S5). Using this method, we determined that our DPAP3 stock contained 840 nM of active protease.

### Substrate turnover assay

The substrate library was screened in triplicate at 1 μM substrate and 1 nM DPAP3 in assay buffer. Substrate turnover was measured over 30 min at RT using a SpectroMax M5e plate reader: *λ*_ex_ = 355 nm, *λ*_em_ = 460 nm, emission filter 455 nm, for ACC or AMC (7-amino-4-methylcoumarin) substrates; *λ*_ex_ = 315 nm, *λ*_em_ = 355 nm, emission filter 420 nm, for Phe-Arg-βNA); and *λ*_ex_ = 492 nm, *λ*_em_ = 523 nm, emission filter 520 nm for (PR)_2_Rho. Calibration curves of free βNA (β-napthylamide) and ACC (0-500 nM) under the same assay conditions were performed to convert the turnover rate measured as fluorescent units per second into moles per second. The determine *k*_cat_ and *K*_m_ values, substrate turnover was measured at different substrate concentrations in assay buffer using 1 nM of DPAP3 or 1 nM of bovine CatC. The initial velocities were then fitted with Prism to the Michaelis Menten Eqs. 3 or 4 to obtain accurate *k*_cat_ and *K*_m_ or *k*_cat_/*K*_m_ values, respectively. A minimum of three replicates were performed for each substrate.

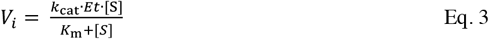

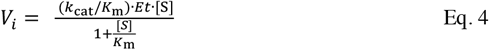

V_i_ is the initial velocity and Et the total concentration of active protease.

### Irreversible inhibition assay

To determine the inhibition constants of vinyl sulfone inhibitors against DPAP3, 2.2 μM of Met-nLeu(o-Bzl)-ACC (0.25 x *K*_m_) was mixed with a dose response of inhibitor in assay buffer, and the turnover rate measured over 40 min at RT after addition of 0.2 nM DPAP3. The data was analyzed according to the irreversible inhibition model shown in Eq. 1. First, the progress curves (fluorescent units vs. time) were fitted to Eq. 5, where F is the measured fluorescence at time t, F_0_ the initial fluorescence, *V*_0_ the initial turnover rate, and *k*_obs_ the observed second order rate constant of inhibition measured at each inhibitor concentration [64].

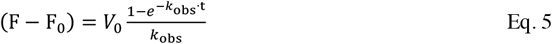

Second, *k*_obs_ values were fitted to Eqs. 6 or 7. to obtain the *k*_inact_ and *K*_i_ or *k*_inact_/*K*_i_ values, respectively.

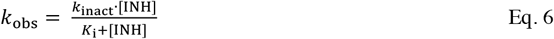

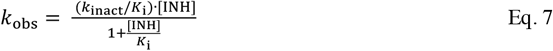

Inhibition constants for DPAP1 and CatC were determined in a similar way but using 0.25 x *K*_m_ concentrations of the Pro-Arg-AMC (20 μM) and Gly-Arg-AMC (15 μM), respectively. Inhibition of DPAP1 was directly measured in parasite lysates diluted 100-fold in assay buffer using the DPAP1-selective substrate Pro-Arg-AMC[41]; 0.2 nM of CatC was used for these inhibition studies.

### Measurement of inhibitor specificity in biological samples

Inhibition of *Plasmodium* DPAPs and the falcipains in intact parasites was measured using the FY01 ABP in competition assays as previously described[25,26]. Because DPAP3 is maximally expressed in very mature schizonts[6], labelling was performed after treating parasites with 1 μM of Compound 2, a cGMP-dependent protein kinase inhibitor that arrest parasite development 15-30 min before they egress from the infected RBC[65]. For each sample, 5 μL of percoll-purified schizonts were diluted in 45 μL of RPMI (Gibco), pre-treated for 30 min with a dose response of inhibitor, and labelled for 1 h with 1 μM FY01. Samples were then boiled for 10 min in loading buffer and run on a 12 % SDS-PAGE gel. Fluorescently labelled proteases in the gel were detected using a Bio-Rad PharosFX flatbed fluorescence scanner.

## Acknowledgement

We would like to thank the Wellcome Trust and Royal Society for funding this research through the Sir Henry Dale Fellowship 099950. We would also like to thank Erasmus for funding L.E.d.V. and L.K. internship in Dr. Edgar Deu’s laboratory. M.M. and M.H. were supported by project ChemBioDrug CZ.02.1.01/0.0/0.0/16_019/0000729 from the European Regional Development Fund (OP RDE) and by the institutional project RVO 61388963.

## Author Contributions

E.D. planned, performed, and managed experiments, analysed data and wrote the manuscript. L.E.d.V., L.K., and C.L. performed experiments. M.I.S., F.Y., S.A.K., M.P., K.G. synthesized compounds under the pupervision of M.B. and M.D. M.H. and M.M. contributed reagents. All authors revised the manuscript.

